# *Status epilepticus* during neurodevelopment increases seizure susceptibility: a model study in zebrafish (*Danio rerio*)

**DOI:** 10.1101/2023.12.20.572588

**Authors:** Samara Cristina Mazon, Ellen Jaqueline Mendes, Adrieli Sachett, Kanandra Taisa Bertoncello, Isabella Bodanese Marsaro, Carla Fatima Daniel, Fernanda Barros de Miranda, Mariana Appel Hort, J. Vladimir Oliveira, Angelo Piato, Cássia Alves Lima-Rezende, Ana Paula Herrmann, Liz Girardi Müller, Anna Maria Siebel

## Abstract

Epilepsy is among the most common neurological diseases, affecting more than 50 million people worldwide. Unfortunately, one-third of people with epilepsy fail to respond to any current treatment and present a decreased quality of life. Nevertheless, different studies suggest that epilepsies acquired from an initial insult as *status epilepticus* (SE) followed by *epileptogenesis* can be prevented. Therefore, it is necessary to establish animal models of *epileptogenesis* induced by SE. Thus, here we proposed an animal model of *epileptogenesis* triggered by SE that occurred during the neurodevelopment of zebrafish (*Danio rerio*). Zebrafish larvae at the 7^th^ day post-fertilization (dpf) were randomly assigned to two experimental groups. One group was exposed to embryo medium. The other group was exposed to pentylenetetrazol (15 mM PTZ) to induce SE. At the 8^th^ dpf, each larva was subjected to the open tank test (OTT) to analyze locomotion and behavioral parameters. After the OTT, each initial group was divided into two groups: animals maintained in embryo medium and animals exposed to PTZ (3 mM), and the susceptibility to PTZ-induced seizure-like behavior was analyzed. Data showed that animals submitted to SE on the 7^th^ dpf showed altered locomotion and behavior 24 hours later. Interestingly, zebrafish larvae submitted to SE on the 7^th^ dpf showed decreased latency to reach seizure-like behavioral stages when exposed to a low concentration of PTZ on the 8^th^ dpf. Concluding, our results show that SE during neurodevelopment increases the susceptibility of zebrafish larvae to present PTZ-induced seizure-like behavior. Our study contributes to the establishment of a model of *epileptogenesis* in developing zebrafish. Finally, the next step is to improve this model by characterizing molecular, biochemical and physiological markers.

## Introduction

According to the World Health Organization (WHO), epilepsy (characterized by the people’s susceptibility to suffering seizures) is among the main chronic neurological disorders, affecting more than 50 million people worldwide (World Health Organization, 2023). Epilepsy is mainly treated through the administration of antiseizure drugs (Zuberi et al., 2022). However, around 30% of people with epilepsy do not respond to available treatments (Zuberi et al., 2022). Thus, these people suffer recurrent seizures, which trigger different comorbidities such as anxiety, depression, and intellectual disability, impairing their life quality (Devinsky et al., 2018).

It is estimated that 25% of epilepsy cases can be prevented. This 25% corresponds to epilepsies acquired from an initial insult such as *status epilepticus* (SE) followed by *epileptogenesis* (World Health Organization, 2023). SE corresponds to a prolonged seizure or two or more consecutive seizures without intervals (Singh et al., 2020). Today, SE is one of the main pediatric emergencies and is one of the main inducers of *epileptogenesis* (Singh et al., 2020). *Epileptogenesis* is a long and progressive process resulting from neuronal insults like SE (Singh et al., 2020). *Epileptogenesis* involves molecular and cellular changes in the brain that culminate in increased susceptibility to seizures, which characterizes epilepsy (Singh et al., 2020).

The molecular and cellular processes involved in *epileptogenesis* triggered by SE are not fully understood, but it is known that neuroinflammation plays a key role in *epileptogenesis*. Studies in animal models shown that there is the participation of inflammatory mediators such as cytokines 1β (IL-1β) and 6 (IL-6), tumor necrosis factor-α (TNF-α), and brain-derived neurotrophic factor (BDNF) (Dey et al., 2016). The molecular and cellular processes occurring in *epileptogenesis* promoted by an initial insult as SE may represent targets for epilepsy prophylaxis for numerous patients (Dey et al., 2016; Vitaliti et al., 2019). To investigate that it is necessary to establish animal models of *epileptogenesis* induced by SE that occurred during neurodevelopment.

Zebrafish (*Danio rerio*) has been widely used in studies that investigate the neurobiological mechanisms related to epilepsy (Baraban, 2021; Budaszewski Pinto et al., 2021; Bertoncello & Bonan, 2023). Also, zebrafish have been used in the research and development of new antiseizure drugs (Baraban, 2021). It is known that seizures in zebrafish cause behavioral, molecular, and electrographic changes similar to those observed in rodents (Baraban, 2021; Siebel et al., 2011). SE induced by kainic acid causes neurochemical and behavioral changes similar to those observed in rodents (Mussulini et al., 2018). Finally, severe seizures induce short-term biochemical changes that reflect an altered long-term behavior (Budaszewski Pinto et al., 2021).

Considering that (i) epilepsy affects more than 50 million people worldwide, (ii) many patients develop epilepsy due to *epileptogenesis* triggered by SE that occurs during neurodevelopment, (iii) epileptogenesis process can represent targets for epilepsy prophylaxis and (iv) zebrafish is a consolidated model in epilepsy research, we proposed the development of an animal model of *epileptogenesis* triggered by SE occurred during neurodevelopment of zebrafish.

## Material and methods

### Animals

Zebrafish larvae were obtained from the breeding of adult zebrafish (*Danio rerio*, wild-type) (3 months), maintained in our institutional bioterium. For the breeding, females and males (2:1) were placed in a breeding tank. The obtained eggs were collected and classified as fertilized and unfertilized. The fertilized eggs were disposed of in plates containing embryo medium (reverse osmosis water reconstituted with Ocean-Tech/Reef Active marine salt, pH 7.00, renewed every day) and maintained in the biochemical oxygen demand incubator (BOD) (26 ± 2 ºC), under a 14/10h light/dark cycle until the 7^th^ day post-fertilization (dpf) (Spence et al., 2008).

All experimental practices respected the National Council for Control of Animal Experimentation recommendations (CONCEA, 2018) and the European Community instructions. Institutional Ethics Committee for Animal Use approved the experimental protocol (CEUA Unochapecó Protocol, #031/2019).

### Experimental design

The experiments were performed according to Figure 1. All analyses were carried out following blinding standards.

**Figure 1.**
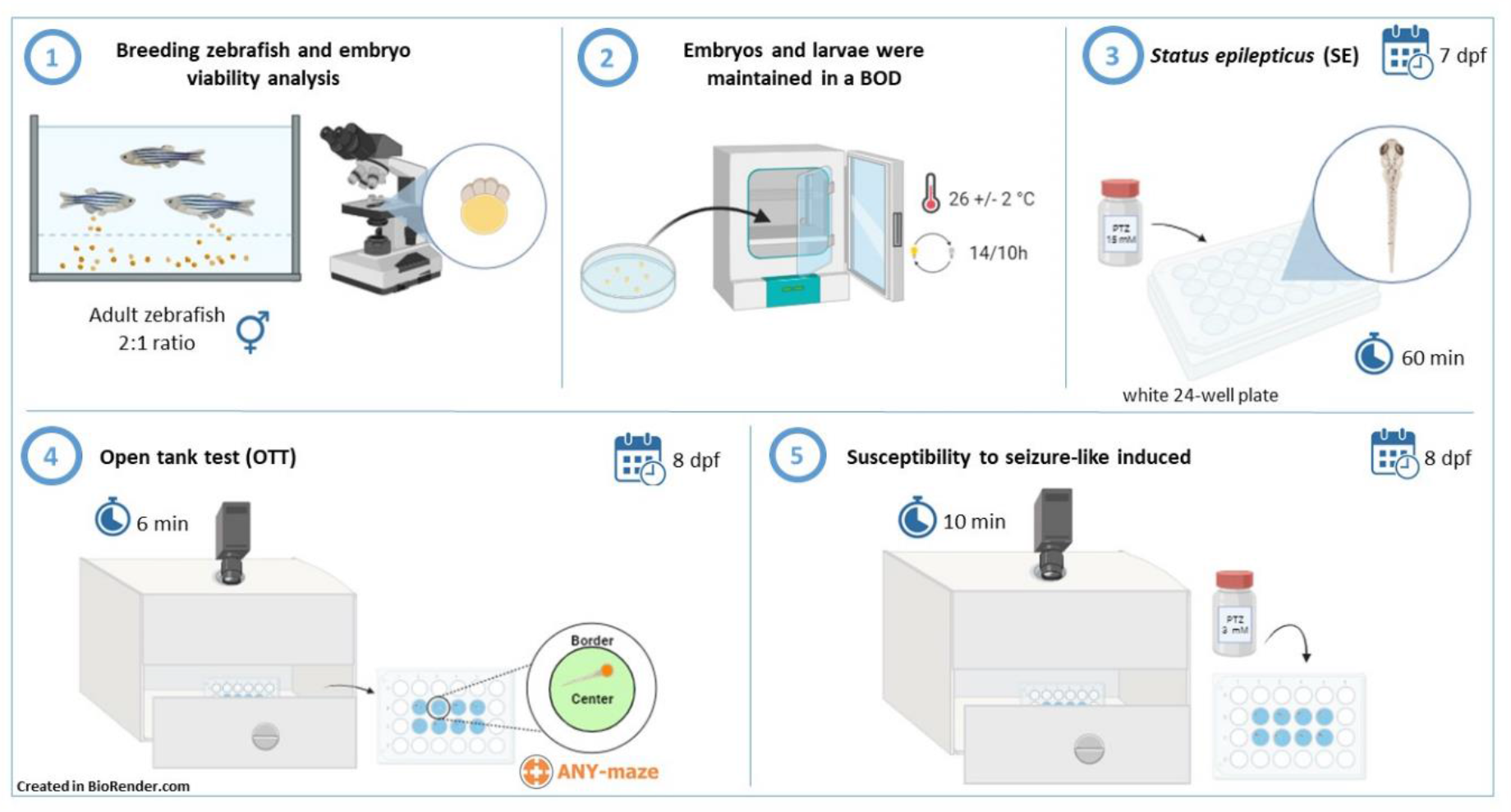
Experimental design. (1) For breeding, the animals were separated at a ratio of two females to each male. Eggs were collected and classified. (2) The embryos and larvae were kept in petri dishes containing embryo medium in BOD. (3) At the 7^th^ dpf, the animals were randomly allocated to two experimental groups and exposed to PTZ for 60 minutes to induce SE. (4) At the 8^th^ dpf, the animals were submitted to the OTT for 6 minutes. (5) Each initial group was divided into two different groups and the animal’s susceptibility to present PTZ-induced seizure-like behavior was analyzed.

At the 7^th^ dpf, each zebrafish larva was randomly allocated to experimental groups (64 animals per group). One group corresponded to larvae disposed of in embryo medium.

The other group corresponded to larvae submitted to SE by exposure to pentylenetetrazole (15 mM PTZ). After this procedure, larvae were placed in 24-well plates for 24 hours.

At the 8^th^ dpf, zebrafish larvae were submitted to open tank test (OTT). Immediately after the OTT protocol, each initial group was divided into two groups: animals maintained in embryo medium and animals exposed to PTZ (3 mM). Finally, the animal’s susceptibility to present PTZ-induced seizure-like behavior was analyzed.

#### *Status epilepticus* (SE)

SE was induced by exposing the animals to the GABAergic antagonist PTZ. At the 7^th^ dpf, each zebrafish larva was individually immersed in a PTZ solution (15 mM) for 60 minutes (white 24-well plates; 8 larvae per plate; 1 larva per well; 1.5 mL of PTZ solution per well) and were recorded from the top view. SE was confirmed when the animals showed PTZ-induced SE-like behavior for more than 30 minutes (Menezes; Rico; Da Silva, 2014).

### Maintenance of larvae from the 7^th^ dpf to the 8^th^ dpf

At the end of SE protocol, the animals were individually placed in a 24-well plate (white 24-well plates; 8 larvae per plate; 1 larva per well; 1.5 mL of embryo medium per well). The plates were maintained in the BOD until the following analysis.

### Open tank test (OTT)

At the 8^th^ dpf, zebrafish larvae’s locomotor activity and behavior patterns were analyzed. Animals were individually allocated (white 24-well plates; 8 larvae per plate; 1 larva per well; 1.5 mL of embryo medium per well) and had their locomotion and behavior recorded.

The animal’s locomotion and behavior were recorded from the top view (for 6 minutes) and were analyzed using ANY-Maze® software (Stoelting Co., USA). The apparatus (each well) was virtually divided into two areas for video analysis: central area (8.2 mm in diameter) and peripheric area (8 mm in diameter) (Sachett et al., 2022). The parameters analyzed were: total traveled distance (meters), immobility time (seconds), time spent in the central area of the well (seconds) and distance traveled in the central area of the well (meters).

The experimental number in OTT was 59-63. The difference occurred because 4 animals died in the SE group after SE exposure. Also, 1 outlier (defined using the ROUT statistical test and considering total distance traveled as the primary outcome) of each group was removed from the analysis.

### Susceptibility to manifest PTZ-induced seizure-like behavior

After OTT, zebrafish each larva was individually (white 24-well plates; 8 larvae per plate; 1 larva per well; with 1.5 mL embryo medium per well) immersed in a low (subconvulsant) concentration (3 mM) of PTZ for 10 minutes (Bertoncello et al., 2018). The exposure to PTZ was recorded on video.

It was analyzed the latency to reach each seizure-like behavior stage: stage I - increased swimming activity; stage II - swimming behavior in whirlpools, and stage III - lost of posture, when the animal falls to the side and remains immobile for 1-3 seconds (Almeida et al., 2021; Baraban et al., 2005; Bertoncello et al., 2018; Garbinato et al., 2021).

The experimental number in this analysis was 29-32. The difference among groups occurred because some animals died after SE induced at the 7^th^ dpf. The group of animals maintained on embryo medium at the 7^th^ dpf and exposed to embryo medium at the 8^th^ dpf presented n=32. The group of animals maintained on embryo medium at the 7^th^ dpf and exposed to PTZ (3 mM) at the 8^th^ dpf presented n=32.The group of animals submitted to SE at the 7^th^ dpf and exposed to embryo medium at the 8^th^ dpf presented n=29. The group of animals submitted to SE at the 7^th^ dpf and exposed to PTZ (3 mM) at the 8^th^ dpf presented n=31.

### Statistical analysis

The sample size was calculated to detect an effect size of 0.3 with a power of 0.8 and an alpha of 0.05 using G*Power 3.1.9.7 for Windows. The suggested total sample size was 128 animals, 32 per experimental group.

The outliers were defined using the ROUT statistical test using the total distance traveled as the primary outcome and were removed from the analyses. Considering OTT data, the test pointed to 1 outlier for the medium group and 1 outlier for the SE group. Therefore, these animals were removed from the analysis.

The normality was confirmed for all data sets using the Shapiro-Wilk test. To investigate the effects of SE occurrence on locomotion and behavioral parameters, it was used Mann-Whitney test (two-tailed). Data are expressed as median with interquartile range. To investigate the effects of SE occurrence on the latency to reach each seizure stage, data were analyzed by two-way ANOVA and Tukey’s *post hoc* test. Data are expressed as mean ± S.D.

The level of significance was set at p<0.05. The graphs were plotted using GraphPad Prism version 8.0.1 for Windows.

## Results

The occurrence of SE at the 7^th^ dpf (SE group) altered the locomotion and behavior of zebrafish larvae at the 8^th^ dpf in comparison with animals that were not submitted to SE (CTRL group) (Figure 2). No significant effects were observed among the groups for total distance traveled (Figure 2A). However, the occurrence of SE at the 7^th^ dpf significantly increased the immobility time (Figure 2B), and decreased the time spent in the center of the apparatus (Figure 2C) and the distance traveled in the center of the apparatus (Figure 2D).

**Figure 2.**
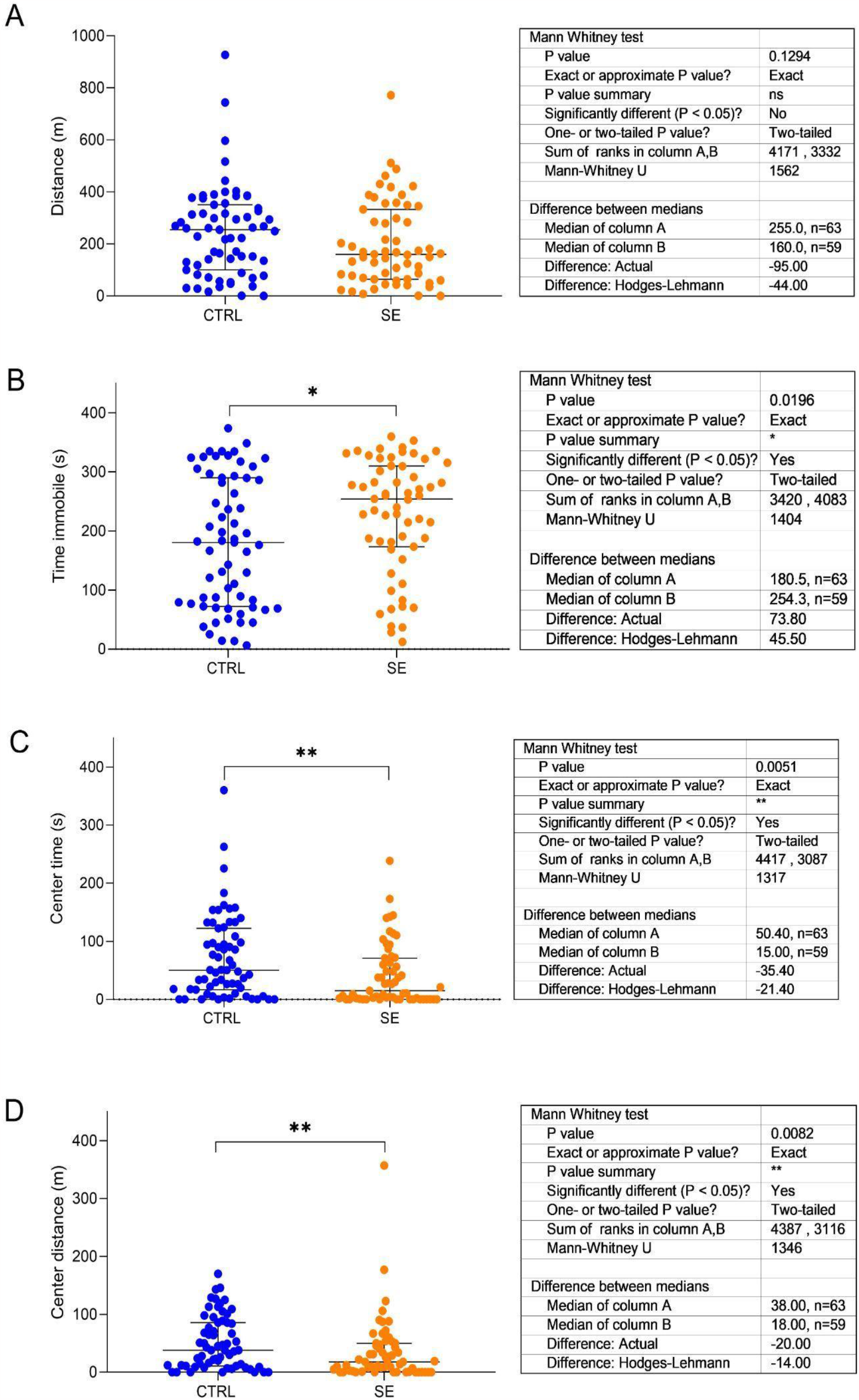
Effects of SE occurred at the 7^th^ dpf on locomotion and behavioral parameters of zebrafish larvae at the 8^th^ dpf. (A) distance, (B) time immobile, (C) center time, (D) center distance. Data are expressed as median with interquartile range. Mann-Whitney test. n= 59-63. *p<0.05, **p<0.01. CTRL vs. SE. *P* and *F* values are presented in the figure.

The effects of the occurrence of SE at the 7^th^ dpf on the susceptibility of zebrafish larvae to present PTZ-induced seizure-like behavior at the 8^th^ dpf are exposed in Figure 3. Zebrafish larvae that were submitted to SE at the 7^th^ dpf showed decreased latency to reach the three seizure stages (Figure 3A, 3B, 3C) when submitted to PTZ (3 mM) at the 8^th^ dpf.

**Figure 3.**
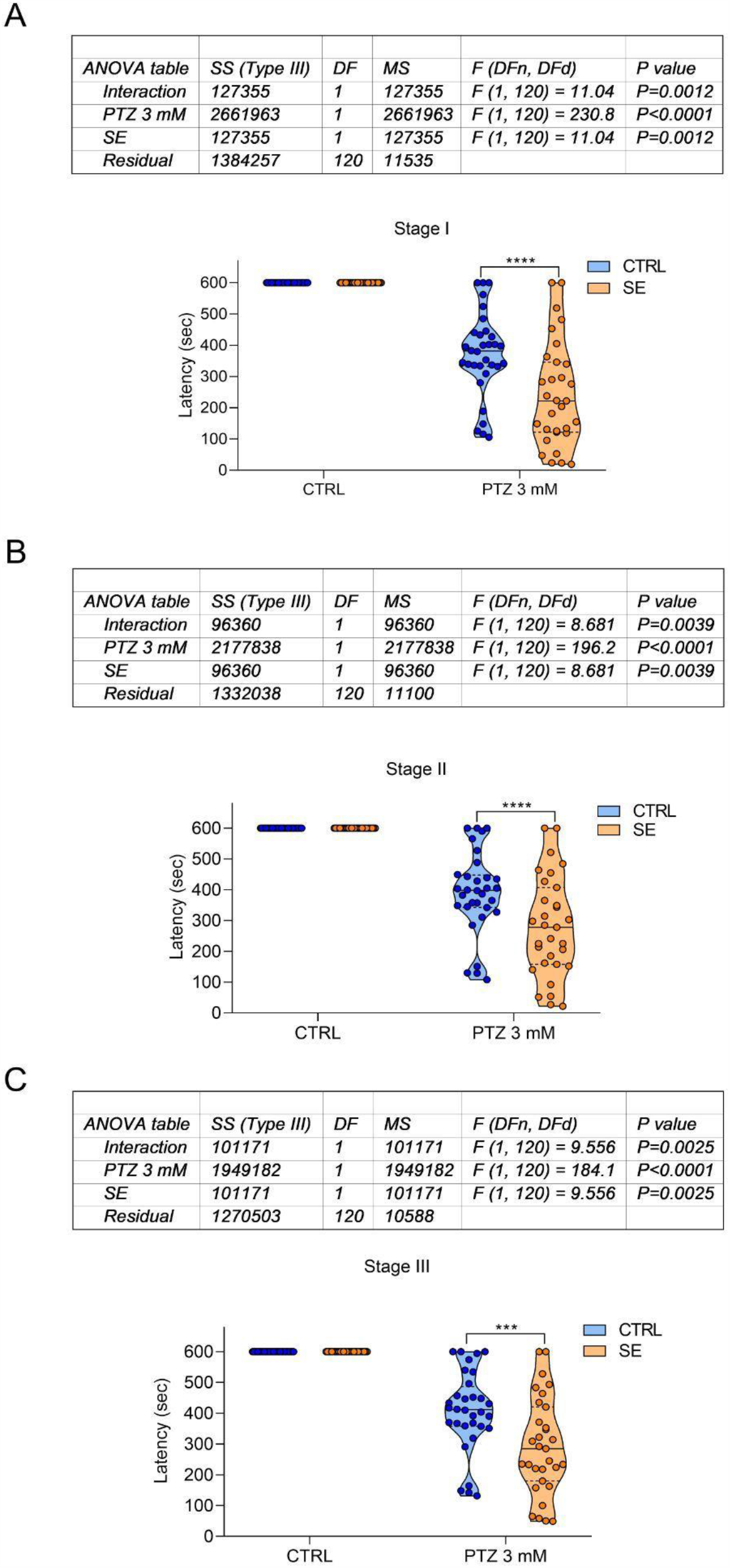
Effects of SE occurred at the 7^th^ dpf on the latency to reach each seizure stage at the 8^th^ dpf. (A) latency to stage I, (B) latency to stage II, and (C) latency to stage III. Data are expressed as mean ± S.D. Two-way ANOVA/Tukey. n = 29-32. ***p<0.001, ****p<0.0001. CTRL vs. CTRL; SE vs. CTRL; CTRL vs. PTZ 3 mM; SE vs. PTZ 3 mM. *P* and *F* values are presented in the figure.

## Discussion and conclusions

Here we found that PTZ-induced SE promoted changes on zebrafish larvae locomotion and behavior: larvae submitted to SE at the 7^th^ dpf increased immobility time and decreased time spent and distance traveled in the center of the apparatus. SE can promote brain damage, causing motor and behavioral changes (Singh et al., 2020). Insults like SE increase the levels of reactive oxygen species on brain cells. This leads to oxidative stress and consequent cell damage and death (Ortiz et al., 2017). Adult zebrafish submitted to pilocarpine-induced seizure-like behavior presented an increased aggressive behavior and increased cell death rate 1 day after pilocarpine exposure (Budaszewski Pinto et al., 2021). We consider that the locomotion and behavioral changes observed here can be associated with cell damage and cell death. However, it is necessary to run additional studies to investigate that. The next steps of this study include to verify possible changes occurring on molecular and biochemical pathways that may explain the results observed in OTT.

Results obtained here showed that zebrafish larvae submitted to SE at 7^th^ dpf decreased the latency to reach the three seizure stages expected for zebrafish larvae when submitted to a subconvulsant concentration of PTZ at the 8^th^ dpf. It is known that pro-inflammatory cytokines are found at low basal levels in the central nervous system and are elevated in response to insults such as SE (Dey et al., 2016). The occurrence of SE in rats caused an increase in the expression of IL-1ß in glial cells for up to 60 days after the episode (De Simoni et al., 2000) IL-1ß potentiates glutamatergic neurotransmission decreasing the astrocytic glutamate up taking (Balosso et al., 2008; Vitaliti et al., 2019). Increased glutamatergic signaling may explain the increased susceptibility to seizures observed in animals submitted to SE (Balosso et al., 2008; Vitaliti et al., 2019). We consider that changes on inflammatory processes can explain the increase on PTZ induced seizure-like susceptibility promoted by SE. Nevertheless, it is necessary to do further studies to investigate this hypothesis.

Taken as a whole, data found here show that SE during neurodevelopment promotes locomotor and behavioral changes and increases seizure susceptibility in zebrafish larvae. The results exposed here represent the first results of our purpose of developing an animal model of *epileptogenesis* triggered by SE that occurred during neurodevelopment. These initial findings will allow the implementation of an innovative animal model for the investigation of new targets for epilepsy prophylaxis, promoting the investigation of new drugs with antiepileptogenic potential.

